# Higher genetic risk for schizophrenia is associated with living in urban and populated areas

**DOI:** 10.1101/179432

**Authors:** Lucia Colodro-Conde, Baptiste Couvy-Duchesne, John B. Whitfield, Fabian Streit, Scott Gordon, Marcella Rietschel, John McGrath, Sarah E. Medland, Nicholas G. Martin

## Abstract

Social stress in urban life has been proposed as an environmental risk factor associated with the increased prevalence of schizophrenia in urban compared to rural areas. However, the potential genetic contributions to this relationship have been largely ignored.

Using a community-based sample of 15,544 adults living in Australia, we found higher genetic loading for schizophrenia in participants living in more densely populated (p-value=5.69*10^-5^) or less remote areas (p-value=0.003). Mendelian Randomization suggested that high schizophrenia genetic risk is a causal factor in chosing to live in denser (p-value=0.046) and less remote areas (p-value=0.044).

Our results support the hypothesis of selective migration to more urban environments by people at higher genetic risk for schizophrenia and suggest a need to refine the social stress model for schizophrenia by including genetic influences on where people choose to live.

---

In 2011, Lederbogen et al., published an influential fMRI study that showed greater brain activation of the stress processing pathways in participants living in urban versus rural areas^1^ and suggested^1,2^ that the greater social stress of urban living could explain could explain the well documented higher prevalence of schizophrenia observed in urban than rural environments (OR =1.72, 95% CI: [1.53-1.92])^3^. Here, we investigate an alternative (but not incompatible) explanation that people with higher genetic risk for schizophrenia tend to live in more urbanized areas due to selective migration^4^ in either past or current generations.

**Supplementary information 1** presents a short review of the literature in this area. In summary, living in an urban environment is associated with increased risk of developing schizophrenia after controlling for potential confounders (age, sex, ethnicity, drug use, social class, family history, season of birth) and using different measures of urbanicity (population size or density^5,6^), window of exposure (birth^4,7^, upbringing^1,5^, or illness onset^6,8^), and disease definition (narrow schizophrenia or broad psychosis^6,9^). While the association is established, its putative (familial) environmental or genetic components are unclear. Although exposure to urban environments has been estimated to account for more than 30% of all schizophrenia cases^9^, it is not clear whether this reflects a causal effect of urban residence on mental health or whether it is a consequence of the disease (i.e. migration to the city of people in the prodromal stages of the disorder^5^).

In the present study we sought to examine the cause of this association by testing whether adults with higher genetic risk for schizophrenia are more likely to live in urbanised and populated areas than those with lower risk. This has been made possible by the advances in the identification of common genetic variants associated with schizophrenia^10^ which have enabled us to calculate the polygenic risk scores (PRS) for schizophrenia in a large community-based sample from Australia for whom genome-wide genotyping is available. In addition, we checked that the association could not be explained by differences in socio-economic status (SES) of the residential areas. We also investigated the direction of causation between schizophrenia and population density using multi-instrument Mendelian randomization. For completeness, we also present the estimates of the heritability and a genome-wide association analysis (GWAS) of our main phenotypes.

## Results

**Supplementary Figure 1** presents the distribution of the variables used in this study. Mean population density was 1,169 people/km^2^ (SD = 1,350, range = 0.01 -5,506). Mean remoteness score was 1.5 (SD=0.72, range: 1-3.6) and mean SES score 6.3 (SD=2.8, range: 1-10). Population density, remoteness and SES were all significantly correlated (**Table 1**).

**Table 1:** Phenotypic correlations (95% CI and p-values) between the demographic variables

|  | Remoteness | SES |
| --- | --- | --- |
| Population density | -0.40 (-0.50 -0.32) | 0.25 (0.16 0.33) |
| Remoteness |  | -0.30 (-0.38 -0.22) |

Phenotypic correlations were calculated in OpenMx <sup>47</sup>, which allows modelling the sample relatedness and shared environment. All correlations were significant (p<0.001) after correcting for multiple testing.

**Figure 1.**
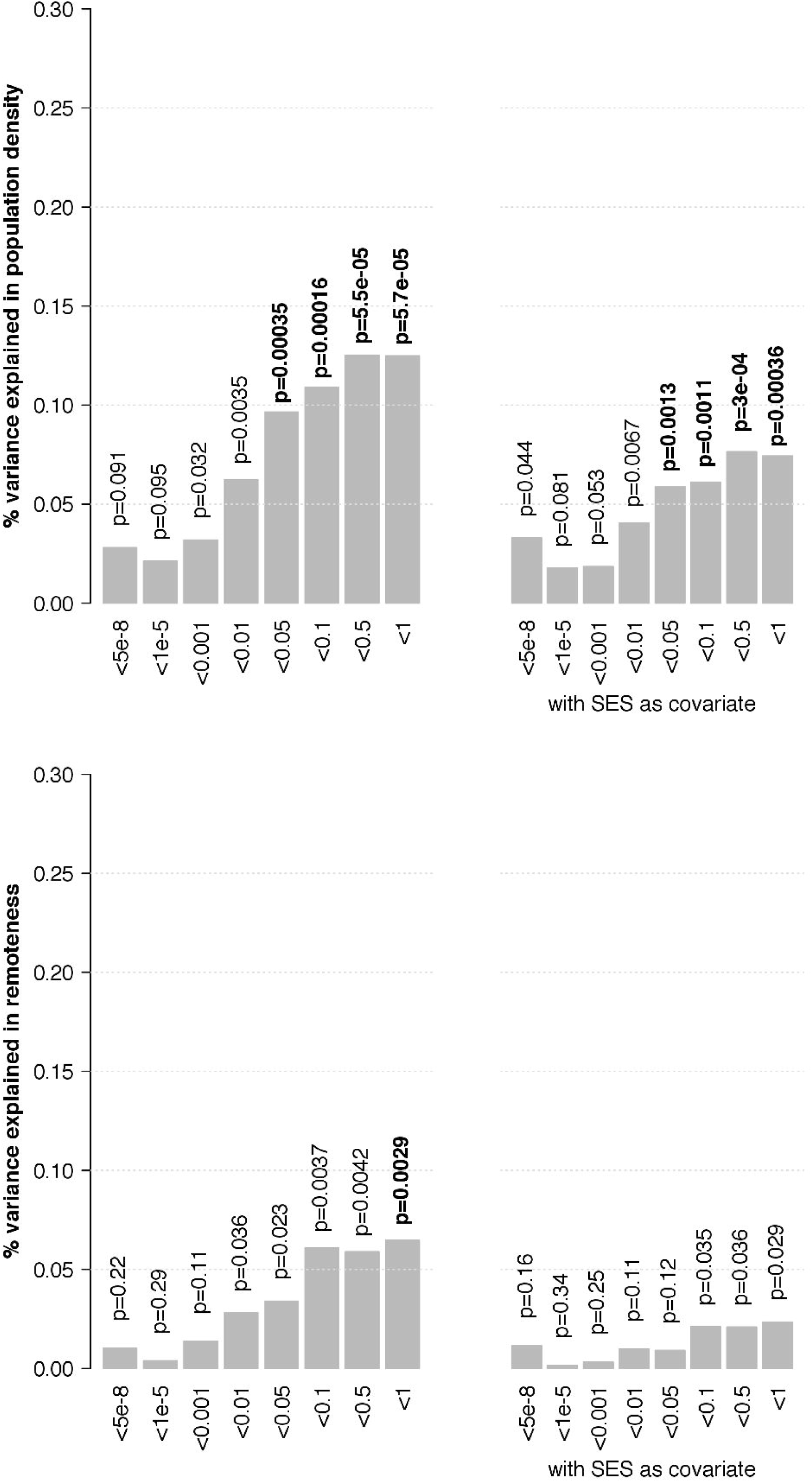
Variance of (a) the population density (b) and remoteness of the place of residence explained by the genetic risk for schizophrenia, both without (left panel) and with (right panel) SES as a covariate.

We first calculated the heritability of our phenotypes using correlations of the 1,119 MZ and 1,104 DZ twin pairs in our sample. Population density and remoteness were marginally heritable (**Supplementary Figure 2**, h^2^_popdensity_ = 16.9%, 95% CI: [3.4-30.4], p-value=0.014, h^2^_remoteness_=16.3% [3.5-29.0] p-value=0.012), while the heritability of SES was not significant (h^2^=11.0% [0.00-24.6], p=0.12), shared environment effects explained a more substantial and highly significant proportion of the trait variance (**Supplementary Figure 2**, c^2^_popdensity_=24.3% [13.1-35.1], c^2^_remoteness_=29.1% [19.0-40.0], c^2^_SES_=26.8% [15.6-37.1], all p-value < 0.001), which highlights that people tend to live with or close to their parents or relatives.

**Figure 2.**
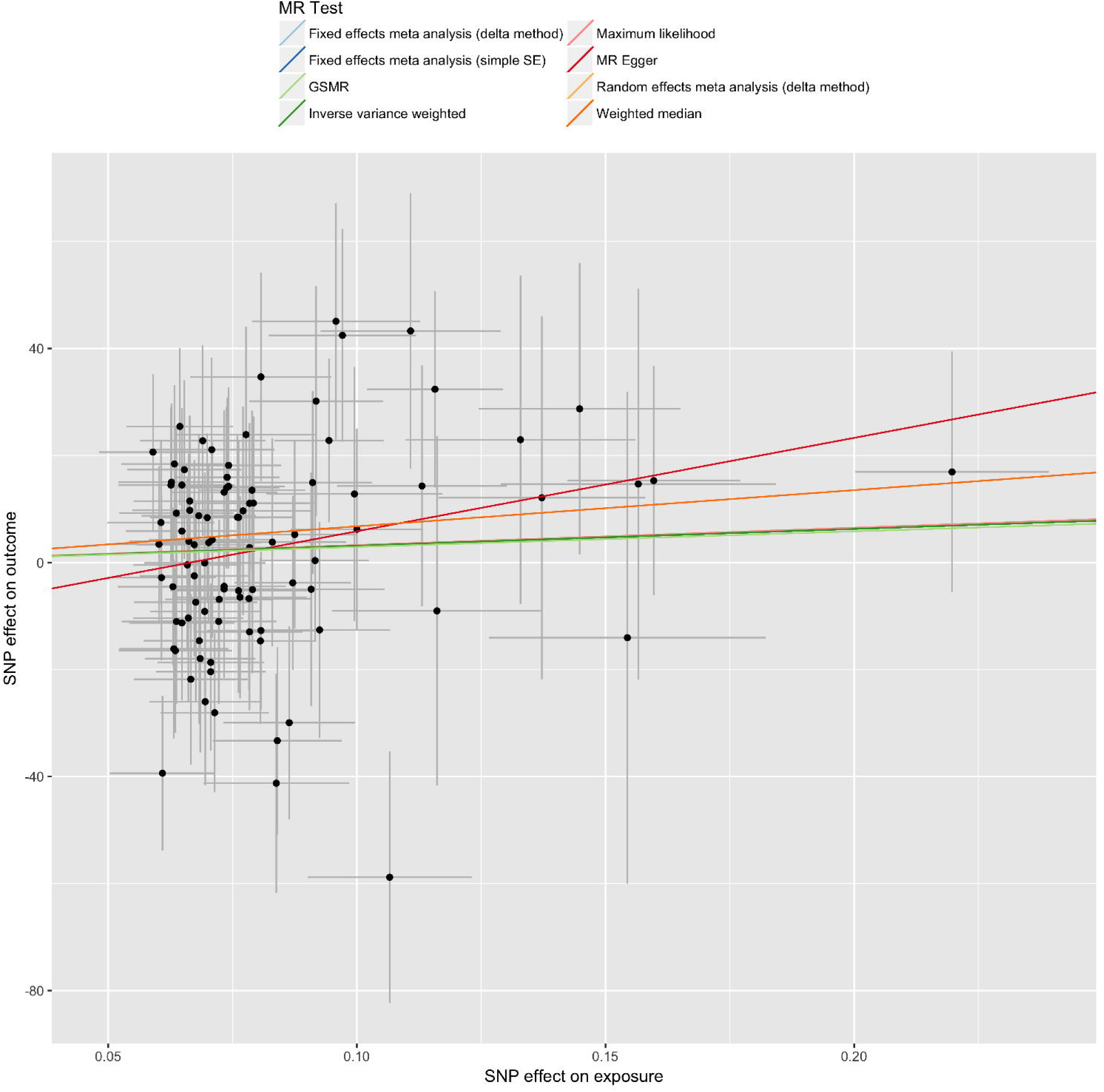
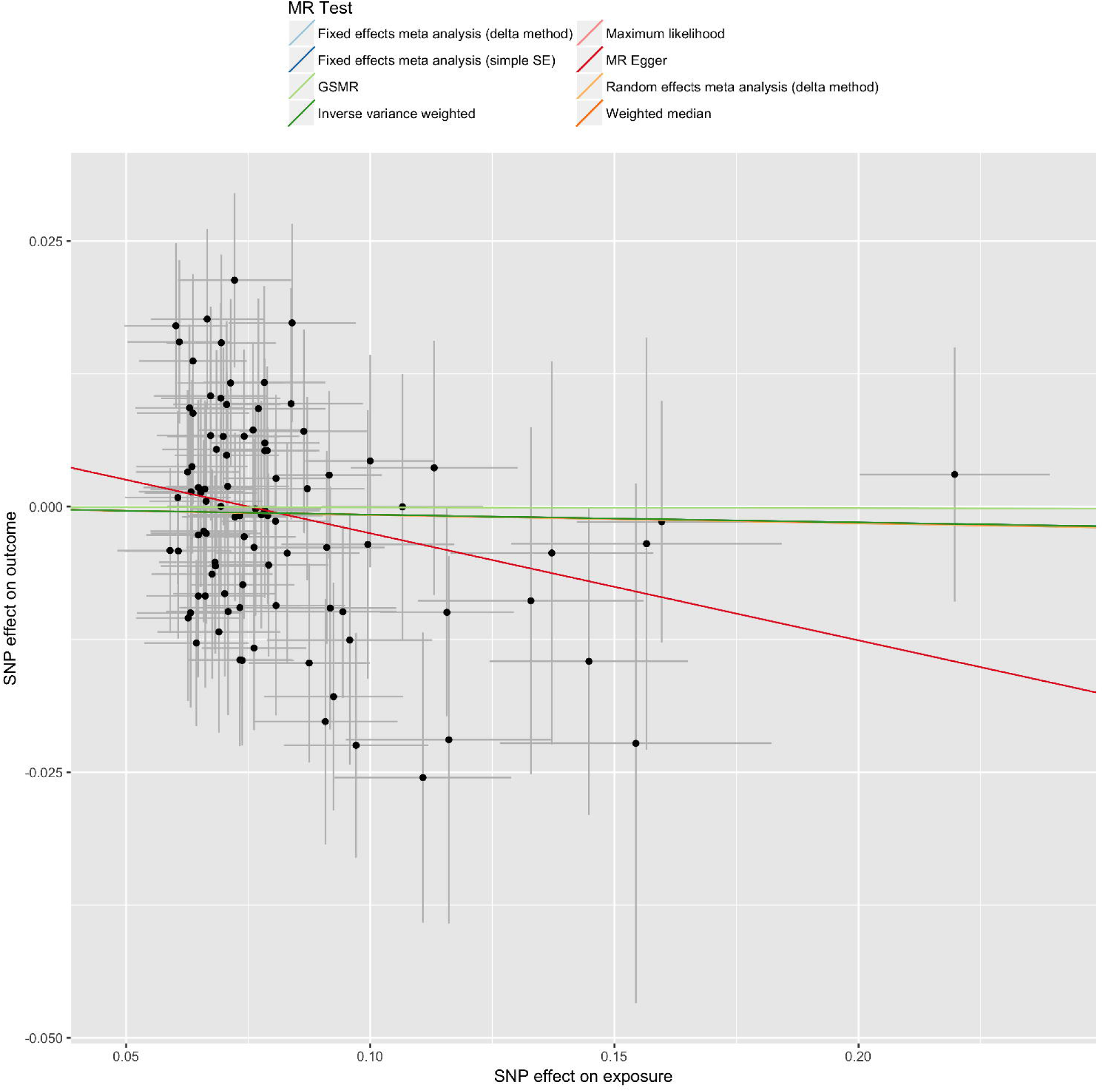
Mendelian Randomization summary results, testing the hypothesis that schizophrenia influences (a) population density and (b) remoteness of place of residence.

We observed an effect of age on the genetic and shared environment sources of variances for population density. The trait was more heritable and less influenced by shared environmental sources as participants got older (**Supplementary Figure 3**) heritability increasing from 9.0 to 25.6% between ages 20 and 80. Over the same life span, C^2^ decreases from 44.5% to less than 10% (**Supplementary Figure 3**), while variance explained by unique environmental sources (including measurement error) remains constant. Similar results were obtained using standardised estimates, suggesting constant phenotypic variance across age. In addition, population density, remoteness and SES shared genetic and environmental influences as indicated by significant genetic and common environmental correlations from the twin models (**Supplementary Table 1**).

**Figure 1a** shows the variance of the population density of the place of residence that is explained by the PRS for schizophrenia, both with and without SES as a covariate; this ignores the potential problem that SES is a genetic covariate of our dependent variable. PRS calculated from all the independent SNPs in the genome (“<1”) explained the greatest amount of variance in population density: r^2^=0.12%, p-value=5.69*10^-5^, and still explained r^2^=0.07% (p-value= 0.0004) when accounting for SES. Schizophrenia PRS also significantly predicted remoteness (r^2^ = 0.06%, p=0.0029) when including all the independent haplotypes (“p-value <1”), although the association disappeared when correcting for SES (**Figure 1b**). The association between SES and schizophrenia PRS did not reach significance after multiple testing correction (p-value>0.013, **Supplementary Figure 4**). The interactions between sex or age and PRS for schizophrenia (p-values > 0.05) that may have accounted for more variance in population density were not significant From the GWAS of population density, no SNP passed the genome-wide significance threshold, regardless of the inclusion of SES as a covariate in the model. The variants presenting the strongest association were located on chromosome 5 (rs79174917, p-value=9.3*10^-8^) or on chromosome 1 (rs144823, p-value=8.3*10^-7^) when including SES as a covariate.

GWAS for remoteness showed a genome-wide significant hit on chromosome 20 (rs7269466, p-value=3.8*10^-8^) (**Supplementary Figure 5**), which disappeared when including SES as a covariate (most significant SNP was rs4691169 in chromosome 4, p-value=1.1*10^-7^). To confirm the results were not due to unaccounted for population stratification, we repeated the GWAS including 20 ancestry principal components. The results were consistent, with the strongest association with remoteness observed in the same SNP in chromosome 20 (rs7269466, p-value=3.9*10^-8^).

Lastly, Mendelian randomization in MR-base^11^ selected 94 genome-wide significant SNPs for schizophrenia after clumping the GWAS summary statistics. This is consistent with the results reported in the Psychiatric Genomics Consortium publication (108 independent associations)^12^; the difference arose from SNPs not included or not passing QC in our GWAS of population density and remoteness. For the MR using population density, there was no evidence of a confounding effect from the heterogeneity of effect sizes (p-value > 0.37) or from pleiotropy (p-value=0.11). The multi-instrument Mendelian randomization only reached significance using the median weighted approach (p-value=0.046; p=0.065 with MR-Egger) (**Table 2, Figure 2a**). Correcting for SES in the GWAS analysis of population density slightly changed the results (p≥0.053, obtained with MR-Egger). The MR using remoteness as outcome showed similar results (**Table 2, Figure 2b**), with no evidence for a confounding effect from heterogeneity of effect sizes (p-value>0.30) or from pleiotropy (p-value=0.05).

**Table 2:**
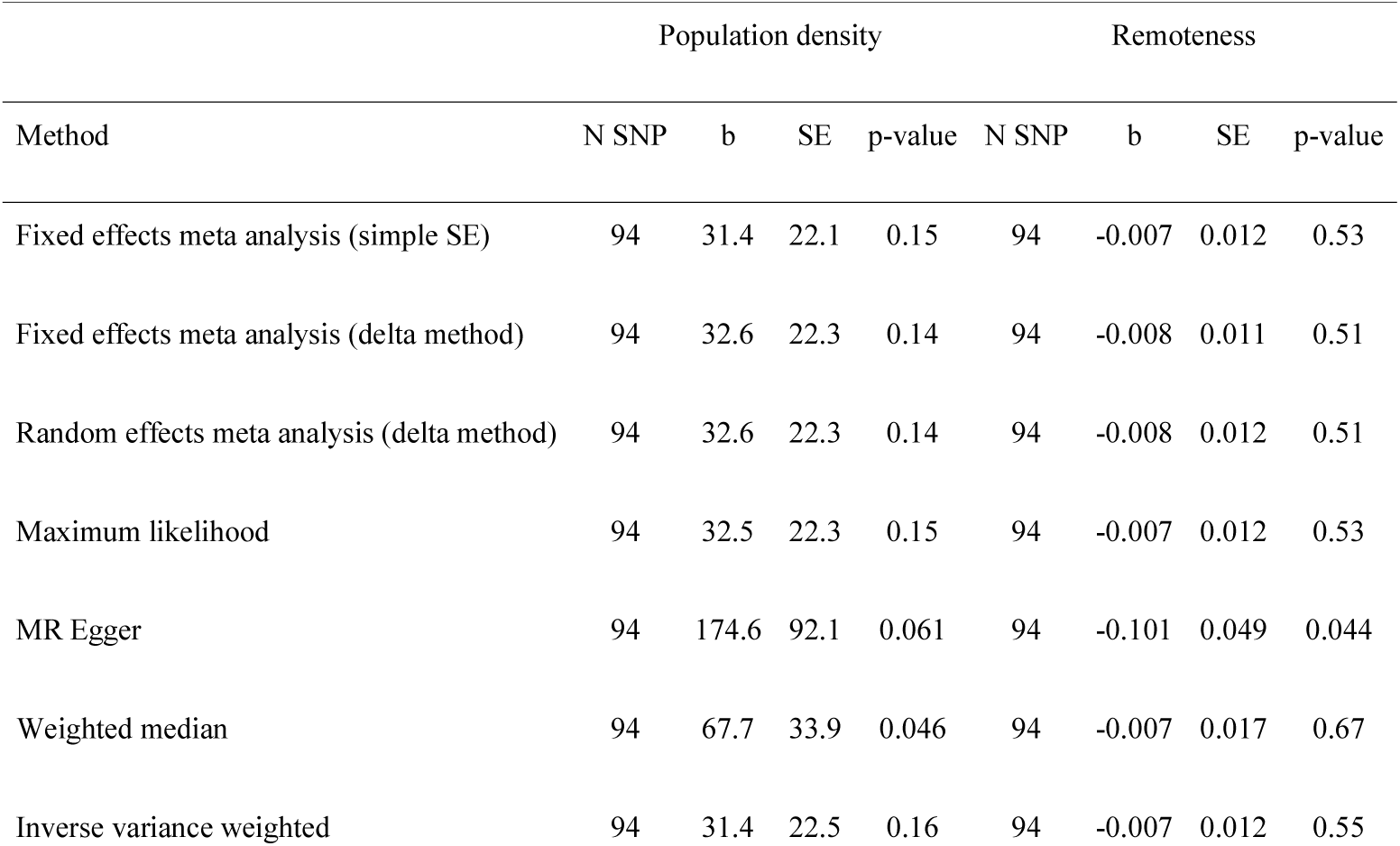
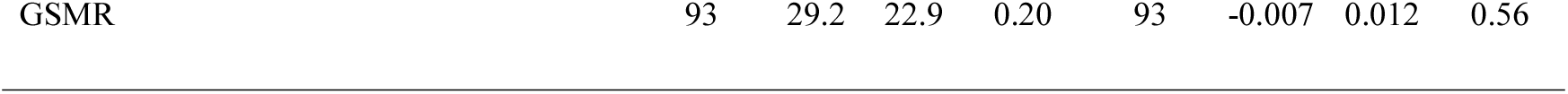
Summary of the MR-base analysis between schizophrenia and population density (left) and remoteness (right), testing the hypothesis that schizophrenia influences population density and remoteness of place of residence.

| Method | Population density |  |  |  | Remoteness |  |  |  |
| --- | --- | --- | --- | --- | --- | --- | --- | --- |
|  | N SNP | b | SE | p-value | N SNP | b | SE | p-value |
| Fixed effects meta analysis (simple SE) | 94 | 31.4 | 22.1 | 0.15 | 94 | -0.007 | 0.012 | 0.53 |
| Fixed effects meta analysis (delta method) | 94 | 32.6 | 22.3 | 0.14 | 94 | -0.008 | 0.011 | 0.51 |
| Random effects meta analysis (delta method) | 94 | 32.6 | 22.3 | 0.14 | 94 | -0.008 | 0.012 | 0.51 |
| Maximum likelihood | 94 | 32.5 | 22.3 | 0.15 | 94 | -0.007 | 0.012 | 0.53 |
| MR Egger | 94 | 174.6 | 92.1 | 0.061 | 94 | -0.101 | 0.049 | 0.044 |
| Weighted median | 94 | 67.7 | 33.9 | 0.046 | 94 | -0.007 | 0.017 | 0.67 |
| Inverse variance weighted | 94 | 31.4 | 22.5 | 0.16 | 94 | -0.007 | 0.012 | 0.55 |
| GSMR | 93 | 29.2 | 22.9 | 0.20 | 93 | -0.007 | 0.012 | 0.56 |

The results were significant using the MR Egger method: p-value=0.044. For both phenotypes, GSMR^13^ retained 93 SNPs after checking for pleiotropy (HEIDI test, p-value>0.01). The causality estimates were in the same direction as those from MR base but did not reach significance (p-value=0.20 for population density, p-value=0.56 for remoteness, Table 2). Our results point to a positive causation for population density and negative for remoteness, such that genetic risk for schizophrenia is a causal factor in the choice to live in more densily populated and less remote areas.

## Discussion

The present study investigated the association between genetic risk for schizophrenia, and living environment (population density, remoteness and SES) in order to test the genetic nature of the relationship between schizophrenia and population density. We used data on where people live collected from a community sample of 15,544 adult Australians with available genetic array information.

Genetic risk for schizophrenia was associated with greater level of urbanicity measured as the population density of the postcode of residence. The strongest association was found with the schizophrenia PRS calculated on all independent SNPs (“< 1”, r^2^=0.12%, p-value=5.69*10^-5^, **Figure 1a**) which reduced to 0.07% when partialling out SES (p-value=0.0004). Our results show that the geographical distribution of the genetic risk for schizophrenia is not uniform and that participants with higher genetic risk levels live in areas with higher population density over what it is expected by chance.

Our Mendelian randomization analysis was suggestive of schizophrenia risk being a causal factor in the choice to live in more densely (positive causation, p-value=0.046) and less remote (negative causation, p-value=0.044) areas, although more powered analyses are required to confirm this results, as well as to investigate the reverse causation. Larger GWAS in psychiatry and of living environment variables may shed light on how the two relate to each other. These results support the selective migration hypothesis, that individuals with genetic liability for schizophrenia tend to move to urban areas^14^. More data is needed to clarify the impact of comorbid psychiatric risks^15^,^16^ and associated traits (e.g. educational attainment, creativity, risk taking)^16-18^ on the reported association.

Our work builds on previous research from our group that reports that density of population of where you live is significantly heritable^19^, as well as evidence of a familial effect (i.e. due to genetics and/or family environment), according to non-molecular studies, in the relationship between schizophrenia and urban dwelling^4,20,21^. Our results complement two recent publications on the interplay between schizophrenia risk and living environment. The first, from the Swedish registries reported an association between schizophrenia PRS and deprived neighbourhood ^22^, which did not replicate in our sample using SES (r^2^_=_0.05%, p-value>0.01); however, our SES measure^23^ may differ from the Swedish study in term of the factors used in its calculation ^22^. Another study from the Danish registries found a non conclusive association between urban living at 15 years old (but not at birth) and schizophrenia PRS^24^. Thus, there may also be a genetic relationship between upbringing environment and the disease risk i.e. a passive gene-environment correlation, where the association is driven by the genotype a child inherits from their parents and the environment in which he/she is raised. Here, we rather focused on active gene-environment correlation, presumably driven by selective migration, by including only older participants who have a higher degree of independence in choosing where they would rather live. More work is needed to confirm and examine these results over age groups, which will likely require large longitudinal cohorts such as national registries.

We highlighted the importance of age in our analysis by replicating and expanding previous results from our group^19^: that place of residence is heritable and the heritability increases over time while the influence of family environment declines. This age effect, together with sex differences in prevalence and age of onset of schizophrenia ^25,26^, justifies the study of interactions between PRS and age and sex that may contribute to the choice of living environment; however, these interactions were not significant in our analyses. Our data confirm the well-known finding that remote or less dense areas tend to exhibit lower levels of wealth as measured by SES, but add the nuance that this is shaped not only by the environment but also by the genotypesof the inhabitants.

Another limitation of our study arising from restricted sample size is that our strongest association is with a PRS that includes all independent SNPs (“<1”), which reflects that the schizophrenia GWAS we drawn on is still underpowered to detect all variants associated with the disease ^27^. This results in a limited PRS instrument that only explains a fraction of the total trait heritability ^27^. Larger GWAS will produce stronger instruments that will provide more power to identify living environments correlated with the genetic risk of schizophrenia.

We can evaluate the extent to which our results are directly transferable to other countries. To note, Australia is one of the least densely populated countries (233^rd^ rank out of 244 countries, http://en.wikipedia.org/wiki/List_of_countries_and_territories_by_population_density) with an average of 3 people per km^2^. However, this average hides the fact that most of the population is concentrated in two coastal regions: the South East (around Brisbane, Sydney and Melbourne) and the South West (around Perth), where the population density approaches those of other developed countries (hence the mean population density of our participants greater than 1,000 inhabitant per km^2^). However, the relative availability of space and the greater opportunities of this young country, may leave more freedom to choose where to live than in other developed countries with high(er) levels of urbanicity and greater historical constraints. Thus, this could account for greater heritability of living environment variables in Australia and smaller environmental contributions^28^. In addition, it may impact the effect sizes reported as suggested by Vassos et al^29^: “given the higher variability of exposure to urban environment, in countries with similar levels of development but less aggregation in cities, the effect of urbanicity could be larger”.

In conclusion, our study provides empirical evidence that the finding that schizophrenia is more prevalent in urbanized areas is not only due to the environmental stressors of the city. We show that the distribution of the genetic risk for the disorder is not uniform and concentrates in more populated and urban areas, supporting the idea of an active gene-environment correlation due to selective migration. Previous evidence of an environmental relationship between city-living and schizophrenia risk^1,2^ is not incompatable with our results and reflects that there are genetic as well as environmental risk factors for schizophrenia. Future disease models will need to include both (genetic) selection and environmental effects of urban stress on schizophrenia.

In addition there is a need to address the potential GxE interactions that would arise if genetic variants influencing schizophrenia also influence the choice of a stressful living environment, which would contribute to the interaction between urbanicity and family history of schizophrenia that has been reported in the Danish population^30^. If indeed urban environment accounts for more than 30% of all schizophrenia cases^14^, the effect reported here (r^2^=0.1% of population density explained) may only account for an extra 0.03% (0.3*0.001) of the cases, but this estimate will likely grow as the PRS for schizophrenia becomes more predictive with increasing GWAS sample size^22^. Such diathesis-stress interaction studies utilizing PRS have already been published for depression ^31,32^, but given the lower prevalence of schizophrenia will likely require national registries.

## Online methods

### Participants

Since 1980, a series of studies of general health conditions conducted by the Genetic Epidemiology Unit at QIMR Berghofer Medical Research Institute (QIMR) have collected longitudinal phenotypic data and genotypes from more than 28,000 individuals as part of ongoing twin and family studies based in Australia ^33,34^. For the present study we selected all genotyped participants over 18 years old, that is, a total N of 15,544 individuals in 7,015 families (65.6% females, age mean: 54.4, SD: 13.2). Importantly, this is a community based sample representative of the general population according to a number of sociodemographic characteristics^33^. Participants were not screened for schizophrenia across the whole sample and we did not control for disease(s) status in the analyses.

Participants were genotyped using commercial arrays (Illumina 317K, 370K, 610K, ‘1st generation’, or Core Exome plus Omni-family, ‘2nd generation’ ^35-37^). Genotype data were cleaned (by batch) for call rate (≥95%); MAF (>=1%); Hardy-Weinberg equilibrium (p≥10^-3^; PLINK1. 9 ^38^), GenCall score (≥0.15 per genotype; mean ≥0.7) and standard Illumina filters. Data were checked for pedigree, sex and Mendelian errors and for non-European ancestry (6SD from the PC1 and/or PC2 means of European populations). DNA was imputed from a common SNP set to the 1000 Genomes (Phase 3 Release 5) ‘mixed population’ reference panel (http://www.1000genomes.org) ^39,40^, a strategy that allows genotype data from different arrays to be combined using a set of SNPs common to the first generation genotyping platforms. Imputation was performed on the Michigan Imputation Server ^41^ using the SHAPEIT/minimac Pipeline ^42,43^ and minimac3 ^44^ or in-house (chromosome X only). ‘1st generation’ and ‘2nd generation’ arrays, with 277,690 and 240,297 core observed markers respectively, were imputed separately due to poor overlap between markers and the two were combined after imputation to maximise sample size. A total of 9,411,304 SNPs were available for analysis, after QC.

This study was approved by the QIMR Berghofer Medical Research Institute’s Human Research Ethics Committee and the storage of the data follows national regulations regarding personal data protection. All the participants provided informed consent.

### Measures

Population density and remoteness were the main measures of urbanicity in the present study. Together with socio economic status (SES), these variables were generated from the postcode in the address provided by the participants, updated to the time of last contact (1990-2015).

Population density, remoteness and SES were based on the most recent data published by the Australian Bureau of Statistics (ABS), Australia’s national statistical agency, in the ABS Census of Population and Housing (version 2016 for population density and 2011 for remoteness and SES). The ABS uses the Australian Statistical Geography Standard (ASGS) for the collection and dissemination of geographically classified statistics, which are comparable and spatially integrated. The boundaries of the units of analysis established by the ASGS take into account continuous changes relating to population and infrastructure ^45^. We have linked the postal codes provided by the participants of the present study to the information presented by postal areas or the second level of statistical areas (statistical area 2 or SA2), which represent communities that interact together socially and economically.

Population density was calculated by dividing the estimated resident population by the km^2^ of each statistical area. Remoteness areas are regions in each state and territory, divided on the basis of their relative access to services^46^ into five levels: major cities (1), inner regional (2), outer regional (3), remote (4) and very remote (5)^46^. We averaged the remoteness scores of participants living at the border of two different areas and treated the variable as continuous. Outlying values for both population density and remoteness were winsorized to 3 standard deviations. SES was based on the Index of Relative Socio-economic Advantage and Disadvantage (IRSAD)^23^, which can be used to measure socioeconomic wellbeing in a continuum, from the most disadvantaged areas (low values) to the most advantaged areas (high values). We used deciles of this index, which allowed comparison within postal areas in the same state/territory. Thus, SES as defined in the present study, is a function of the area where a person lives and not of any personal characteristics.

## Statistical analyses

### Heritability analysis of population density, city living and SES

We used the 5,894 twins from the full sample, to estimate the contribution of additive genetic (narrow sense twin heritability), shared/familial environment and unique environment to the inter-personal differences in type of residence. Our sample comprised 1,119 complete MZ pairs of twins, 1,104 complete DZ pairs and 1,448 singleton twins. The twins were 53 years old on average (SD=12.8, range 18-99) and mostly (65%) females. We used the OpenMx ^47^ package in R ^48^ to estimate the parameters of the mixed models. Significance of the variance components was tested using likelihood ratio tests on nested models ^49^.

In addition, we fit a GE moderator effect model ^50^ that allows the variance components (heritability, shared environment and unique environment) to vary across age. This approach reflected previous results from our group that suggested different heritability and shared environment over age on rural/urban living in Australia ^19^. The model is as follows^51^:

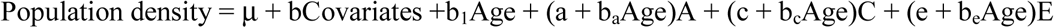

With b and b_1_ the fixed effects parameters of age and other covariates effects on population density mean; a, c, e, b_a,_ b_c,_ and b_e_ the parameters of the random effects. A, C and E are the random effect of additive genetics, shared environment and unique environment assumed to follow a normal distribution of mean 0 and of known variance covariance (kinship matrix) for A, family indicator for C, and identity matrix for E.

Finally, we estimated the genetic and environmental correlations between population density, remoteness and SES using bivariate twin models ^49^. All models included age, age^2^, sex, age*sex, age^2^*sex, GWAS chip and 4 genetic principal components as covariates.

### Polygenic risk scores predictions

Polygenic risk scores ^52,53^ were calculated from the imputed genotype dosages, using GWAS summary statistics from the most recent GWAS meta-analysis from the 2014 Psychiatric Genomics Consortium (PGC) schizophrenia working group ^54^. From our data, we excluded SNPs with low imputation quality (r^2^<0.6) and MAF below 1%. We selected the most significant independent SNPs using PLINK1.9 ^38^ in order to correct for signal inflation due to Linkage Disequilibrium (LD) (criteria LD r^2^<0.1 within windows of 10MBp). We calculated 8 different PRS using different p-value thresholding of the GWAS summary statistics (see **Supplementary Table 2** for number of SNPs included in each threshold and **Supplementary Figure 1** for histograms of PRS for SCZ).

In order to estimate the variance in population density explained by the PRS, we fit linear mixed models which controlled for relatedness and demographic covariates. The parameters of the model were estimated using GCTA 1.26.0 ^55^ (Student’s t-test to test the significance of the fixed effects) that accounts for twin relatedness using a Genetic Relatedness Matrix (GRM). The linear model used is as follows:

Population density or remoteness = intercept + b*Covariates + c*PRS + G

With b, c the vectors of fixed effects

Covariates = age, age^2^, sex*age, sex*age^2^, sex, GWAS array, first 4 genetic principal components, with and without SES.

G is the random effect that models the sample relatedness G∼(0, GRM*σ^2^_G_), with GRM the N*N matrix of relatedness estimated from SNPs (with N the number of individuals).

Formally, we performed 40 tests. However, after taking into account the correlation between the variables^56,57^ (see https://neurogenetics.qimrberghofer.edu.au/SNPSpDlite/ for scripts and details), the number of effective tests is 16. Therefore we used a significance threshold of 3.15 *10^-3^ (Bonferroni correction). This was calculated from an estimated four effective PRS (out of eight used) and four effective phenotypes (out of five considered: population density, remoteness, SES as well as the first two after regressing SES). Such approach represents a fast and efficient alternative to permutation testing^56^, when testing correlated variables.

### Post hoc interaction analysis

When PRS and age or sex were significant, we tested for the presence of interactions. First, we tested for sex specific effects by adding an interaction term (PRS*sex) to the model and using residualized PRS (we regressed out all covariates but sex and its interactions) to remove the confounding effect of the covariates^58^. Similarly, we tested for a PRS by age interaction.

### Genome Wide Association Analyses

Genome-wide association analyses of population density and remoteness were conducted using linear regression under a model of additive allelic effects with age, age^2^, sex, age*sex, age^2^*sex, SES, GWAS array and 4 genetic principal components as covariates using RAREMETALWORKER^59^. We explicitly corrected for relatedness using the –kinpedigree option. Those SNPs with a MAF<0.05% and imputation *r*^2^<0.6 were excluded, leaving 8,495,074 SNPs for analyses.

### Mendelian randomization: is schizophrenia genetic risk influencing where you live or does where you live influence schizophrenia risk?

Here, we relied on the 2014 GWAS summary statistics from the Psychiatric Genetic Consortium (PGC) for schizophrenia and used the GWAS results calculated from our sample for first, population density and second, remoteness. We used MR-base^11^ (“TwoSampleMR” R package^60^) to conduct Mendelian randomization using known schizophrenia SNPs as instruments, thus testing the selection hypothesis of schizophrenia causing to live in a denser and less remote area. Multi instrument Mendelian randomization relates the effect sizes of the exposure SNPs (i.e. genome wide significant SNPs for schizophrenia) to their effect sizes on the outcome (from GWAS of population density and remoteness). The slope is an estimate of the causal effect of the exposure on the outcome. Meta-analysis methods are used to estimate this slope as effect sizes have different precision (from different MAF or different number of observations)^61^. Note that the inference of causation is limited by the instrument SNPs included, that likely only represent a fraction of the biological pathways involved in schizophrenia, due to limited power in GWAS.

First, we merged the GWAS results from the PGC and from our sample, and selected only the genome-wide significant SNPs for schizophrenia. Then, we clumped the SNP list, keeping the most associated SNP (with schizophrenia) per haplotype (default options in MR base: 10MBp window, r^2^<0.01 using 1000genomes as reference). This maximises the number of SNPs, hence the power of the analysis. MR-base provides estimates of Mendelian randomization using several meta-analysis approaches: fixed effects random effects, maximum likelihood, MR Egger, Weighted Median and Inverse Variance Weighted. In addition, it tests if the results may be confounded by pleiotropy (i.e. different SNPs in LD acting on each phenotype) or by a handful of SNPs (heterogeneity test, like in meta-analyses)^11^. MR-Egger is currently seen as the most robust multi-instrument mendelian randomization approach ^61^.

In addition, we used GSMR (Generalised Summary-data-based Mendelian Randomisation)^13^, which is on average more powerful than the MR Egger approach (as it models residual LD between SNPs) and more robust (as it allows testing for pleiotropy at a SNP level: HEIDI test^13^, vs. testing across all the instruments in MR Egger). We included the same list of pruned SNP in the GSMR analysis and excluded instruments that showed significant pleiotropy (HEIDI test<0.01). We calculated the LD matrix between usable SNPs from our sample using PLINK and GCTA (see http://cnsgenomics.com/software/gsmr/ for code and examples).

We could not perform a reverse MR analysis (testing if population density or remoteness cause schizophrenia) because we did not have multiple instrumental variables (SNPs) associated with population density and remoteness.

## Acknowledgments

Phenotype collection, DNA collection and genotyping were funded by NHMRC (981351) and NIH (R01 AA013326, R01 AA007535, R01 AA010249, R01 AA013321, R37 AA007728) grants to NGM over the past three decades. We thank Andrew Heath, Pam Madden and Grant Montgomery for their contributions to funding and executing of genotyping for this study. LCC was supported by a QIMR Berghofer Fellowship. JMG was supported by NHMRC John Cade Fellowship (APP1056929), and a Niels Bohr Professorship from the Danish National Research Foundation. SEM was supported by a NHMRC research fellowship (1103623). We are grateful to the QIMR participants, data collectors and data managers.

## Supplementary information

### *By order of appearance in text

**Supplementary Information 1.** Short review on the gap in schizophrenia rates between urban and rural environments

**Supplementary Figure 1**. Distribution of the population density, remoteness and polygenic risk scores.

**Supplementary Figure 2**. Variance explained by additive genetic (A), shared environmental (C) and unique environmental (E) factors in population density, remoteness and SES.

**Supplementary Figure 3.** Effect of age on the genetic (A) and environmental (common, C and unique, E) sources of variances for population density.

**Supplementary Table 1.** Genetic (above diagonal) and environmental correlations (shared environment followed by unique environment, below diagonal)

**Supplementary Figure 4.** Variance of SES explained by the genetic risk for schizophrenia.

**Supplementary Figure 5.** Manhattan plot (a) and QQ plot (b) for GWAS of remoteness, followed by regional plot (c) of the top hit.

**Supplementary Table 2.** Number of SNPs included in the calculation of each of the polygenic risk scores

## References

1 Lederbogen, F. et al. City living and urban upbringing affect neural social stress processing in humans. Nature 474, 498-501, doi:10.1038/nature10190 (2011).

2 Lederbogen, F., Haddad, L. & Meyer-Lindenberg, A. Urban social stress-risk factor for mental disorders. The case of schizophrenia. Environ Pollut 183, 2-6, doi:10.1016/j.envpol.2013.05.046 (2013).

3 Krabbendam, L. & van Os, J. Schizophrenia and urbanicity: a major environmental influence-conditional on genetic risk. Schizophr Bull 31, 795-799, doi:10.1093/schbul/sbi060 (2005).

4 Mortensen, P. B. et al. Effects of family history and place and season of birth on the risk of schizophrenia. N Engl J Med 340, 603-608, doi:10.1056/NEJM199902253400803 (1999).

5 Pedersen, C. B. & Mortensen, P. B. Evidence of a dose-response relationship between urbanicity during upbringing and schizophrenia risk. Archives of general psychiatry 58, 1039–1046 (2001).

6 Sundquist, K., Frank, G. & Sundquist, J. Urbanisation and incidence of psychosis and depression: follow-up study of 4.4 million women and men in Sweden. The British journal of psychiatry : the journal of mental science 184, 293–298 (2004).

7 Pedersen, C. B. & Mortensen, P. B. Family history, place and season of birth as risk factors for schizophrenia in Denmark: a replication and reanalysis. The British journal of psychiatry : the journal of mental science 179, 46–52 (2001).

8 Marcelis, M., Takei, N. & van Os, J. Urbanization and risk for schizophrenia: does the effect operate before or around the time of illness onset? Psychological medicine 29, 1197–1203 (1999).

9 Marcelis, M., Navarro-Mateu, F., Murray, R., Selten, J. P. & Van Os, J. Urbanization and psychosis: a study of 1942-1978 birth cohorts in The Netherlands. Psychological medicine 28, 871–879 (1998).

10 Owen, M. J., Sawa, A. & Mortensen, P. B. Schizophrenia. Lancet 388, 86-97, doi:10.1016/S0140-6736(15)01121-6 (2016).

11 Hemani, G. et al. MR-Base: a platform for systematic causal inference across the phenome using billions of genetic associations. bioRxiv, doi:10.1101/078972 (2016).

12 Ripke, S. et al. Biological insights from 108 schizophrenia-associated genetic loci. Nature 511, 421-+, doi:10.1038/nature13595 (2014).

13 Zhu, Z. et al. Causal associations between risk factors and common diseases inferred from GWAS summary data. bioRxiv, doi:10.1101/168674 (2017).

14 Sariaslan, A. et al. Schizophrenia and subsequent neighborhood deprivation: revisiting the social drift hypothesis using population, twin and molecular genetic data. Transl Psychiat 6, doi:10.1038/tp.2016.62 (2016).

15 Bulik-Sullivan, B. et al. An atlas of genetic correlations across human diseases and traits. Nat Genet 47, 1236-1241, doi:10.1038/ng.3406 (2015).

16 Anttila, V. et al. Analysis of shared heritability in common disorders of the brain. bioRxiv, doi:10.1101/048991 (2016).

17 Power, R. A. et al. Polygenic risk scores for schizophrenia and bipolar disorder predict creativity. Nat Neurosci 18, 953-+, doi:10.1038/nn.4040 (2015).

18 Maxwell, J. & O’Reilly, P. F. Schizophrenia and autism polygenic risk scores highlight interplay between behaviour and psychiatry.

19 Whitfield, J. B., Zhu, G., Heath, A. C. & Martin, N. G. Choice of residential location: chance, family influences, or genes? Twin Res Hum Genet 8, 22-26, doi:10.1375/1832427053435391 (2005).

20 Cantor-Graae, E. & Selten, J. P. Schizophrenia and migration: a meta-analysis and review. The American journal of psychiatry 162, 12-24, doi:10.1176/appi.ajp.162.1.12 (2005).

21 Pedersen, C. B. & Mortensen, P. B. Are the cause(s) responsible for urban-rural differences in schizophrenia risk rooted in families or in individuals? American journal of epidemiology 163, 971-978, doi:10.1093/aje/kwj169 (2006).

22 Sariaslan, A. et al. Schizophrenia and subsequent neighborhood deprivation: revisiting the social drift hypothesis using population, twin and molecular genetic data. Translational psychiatry 6, e796, doi:10.1038/tp.2016.62 (2016).

23 Australian Boreau of Statistics. Measures of Socioeconomic Status [ABS Catalogue no. 1244.0.55.001]. (2011).

24 Paksarian, D. et al. The role of genetic liability in the association of urbanicity at birth and during upbringing with schizophrenia in Denmark. Psychological medicine, 1-10, doi:10.1017/S0033291717001696 (2017).

25 Abel, K. M., Drake, R. & Goldstein, J. M. Sex differences in schizophrenia. Int Rev Psychiatry 22, 417-428, doi:10.3109/09540261.2010.515205 (2010).

26 McGrath, J. et al. A systematic review of the incidence of schizophrenia: the distribution of rates and the influence of sex, urbanicity, migrant status and methodology. BMC medicine 2, 13, doi:10.1186/1741-7015-2-13 (2004).

27 Dudbridge, F. Power and predictive accuracy of polygenic risk scores. PLoS Genet 9, e1003348, doi:10.1371/journal.pgen.1003348 (2013).

28 Willemsen, G., Posthuma, D. & Boomsma, D. I. Environmental factors determine where the Dutch live: results from the Netherlands twin register. Twin Res Hum Genet 8, 312-317, doi:10.1375/1832427054936655 (2005).

29 Vassos, E., Pedersen, C. B., Murray, R. M., Collier, D. A. & Lewis, C. M. Meta-analysis of the association of urbanicity with schizophrenia. Schizophr Bull 38, 1118-1123, doi:10.1093/schbul/sbs096 (2012).

30 van Os, J., Pedersen, C. B. & Mortensen, P. B. Confirmation of synergy between urbanicity and familial liability in the causation of psychosis. Am J Psychiat 161, 2312-2314, doi:DOI 10.1176/appi.ajp.161.12.2312 (2004).

31 Mullins, N. et al. Polygenic interactions with environmental adversity in the aetiology of major depressive disorder. Psychol Med 46, 759-770, doi:10.1017/S0033291715002172 (2016).

32 Colodro-Conde, L. et al. A direct test of the diathesis-stress model for depression. Mol Psychiatry, doi:10.1038/mp.2017.130 (2017).

33 Heath, A. C. et al. Genetic and environmental contributions to alcohol dependence risk in a national twin sample: consistency of findings in women and men. Psychological medicine 27, 1381–1396 (1997).

34 Knopik, V. S. et al. Genetic effects on alcohol dependence risk: re-evaluating the importance of psychiatric and other heritable risk factors. Psychological medicine 34, 1519–1530 (2004).

35 Cuellar-Partida, G. et al. WNT10A exonic variant increases the risk of keratoconus by decreasing corneal thickness. Human molecular genetics 24, 5060-5068, doi:10.1093/hmg/ddv211 (2015).

36 Medland, S. E. et al. Common variants in the trichohyalin gene are associated with straight hair in Europeans. Am J Hum Genet 85, 750-755, doi:10.1016/j.ajhg.2009.10.009 (2009).

37 McEvoy, B. P. et al. Geographical structure and differential natural selection among North European populations. Genome Res 19, 804-814, doi:10.1101/gr.083394.108 (2009).

38 Purcell, S. et al. PLINK: a tool set for whole-genome association and population-based linkage analyses. Am J Hum Genet 81, 559-575, doi:10.1086/519795 (2007).

39 Genomes Project, C. et al. A global reference for human genetic variation. Nature 526, 68-74, doi:10.1038/nature15393 (2015).

40 Sudmant, P. H. et al. An integrated map of structural variation in 2,504 human genomes. Nature 526, 75-81, doi:10.1038/nature15394 (2015).

41 Whitcher, B., Schmid, V. & Thornton, A. Working with the DICOM and NIfTI Data Standards in R. Journal of Statistical Software 44, 1–28. (2011).

42 Delaneau, O., Marchini, J. & Zagury, J. F. A linear complexity phasing method for thousands of genomes. Nat Methods 9, 179-181, doi:10.1038/nmeth.1785 (2012).

43 Delaneau, O., Zagury, J. F. & Marchini, J. Improved whole-chromosome phasing for disease and population genetic studies. Nat Methods 10, 5-6, doi:10.1038/nmeth.2307 (2013).

44 Howie, B., Fuchsberger, C., Stephens, M., Marchini, J. & Abecasis, G. R. Fast and accurate genotype imputation in genome-wide association studies through pre-phasing. Nature Genetics 44, 955-+, doi:10.1038/ng.2354 (2012).

45 Australian Boreau of Statistics. Australian Statistical Geography Standard (ASGS): Volume 1 - Main Structure and Greater Capital City Statistical Areas [ABS Catalogue No. 1270.0.55.001]. (2011).

46 Australian Boreau of Statistics. Australian Statistical Geography Standard (ASGS): Volume 5 - Remoteness Structure. (2011).

47 Boker, S. et al. OpenMx: An Open Source Extended Structural Equation Modeling Framework. Psychometrika 76, 306–317 (2011).

48 R: A Language and Environment for Statistical Computing (R Foundation for Statistical Computing, Vienna, Austria, 2012).

49 Neale, M. C. & Cardon, L. R. Methodology for genetic studies of twins and families. (Kluwer Academic Pub, 1992).

50 Purcell, S. Variance components models for gene-environment interaction in twin analysis. Twin Res 5, 554-571, doi:10.1375/136905202762342026 (2002).

51 Martin, N. G., Eaves, L. J. & Heath, A. C. Prospects for detecting genotype X environment interactions in twins with breast cancer. Acta Genet Med Gemellol (Roma) 36, 5–20 (1987).

52 Wray, N. R., Goddard, M. E. & Visscher, P. M. Prediction of individual genetic risk of complex disease. Curr Opin Genet Dev 18, 257-263, doi:10.1016/j.gde.2008.07.006 (2008).

53 Wray, N. R. et al. Research Review: Polygenic methods and their application to psychiatric traits. J Child Psychol Psyc 55, 1068-1087, doi:10.1111/jcpp.12295 (2014).

54 Schizophrenia Working Group of the Psychiatric Genomics, C. Biological insights from 108 schizophrenia-associated genetic loci. Nature 511, 421-427, doi:10.1038/nature13595 (2014).

55 Yang, J., Lee, S. H., Goddard, M. E. & Visscher, P. M. GCTA: a tool for genome-wide complex trait analysis. Am J Hum Genet 88, 76-82, doi:10.1016/j.ajhg.2010.11.011 (2011).

56 Li, M. X., Yeung, J. M., Cherny, S. S. & Sham, P. C. Evaluating the effective numbers of independent tests and significant p-value thresholds in commercial genotyping arrays and public imputation reference datasets. Hum Genet 131, 747-756, doi:10.1007/s00439-011-1118-2 (2012).

57 Li, J. & Ji, L. Adjusting multiple testing in multilocus analyses using the eigenvalues of a correlation matrix. Heredity 95, 221–227 (2005).

58 Keller, M. C. Gene x environment interaction studies have not properly controlled for potential confounders: the problem and the (simple) solution. Biol Psychiatry 75, 18–24, doi:10.1016/j.biopsych.2013.09.006 (2014).

59 Sullivan, P. F., Kendler, K. S. & Neale, M. C. Schizophrenia as a complex trait: evidence from a meta-analysis of twin studies. Archives of general psychiatry 60, 1187-1192, doi:10.1001/archpsyc.60.12.1187 (2003).

60 Hemani, G., Haycock, P. & Zheng, J. MR-Base: a platform for systematic causal inference across the phenome using billions of genetic associations. bioRxiv. doi:https://doi.org/10.1101/078972 (2017).

61 Bowden, J., Smith, G. D. & Burgess, S. Mendelian randomization with invalid instruments: effect estimation and bias detection through Egger regression. Int J Epidemiol 44, 512-525, doi:10.1093/ije/dyv080 (2015).

